# Silicon Nanowires for Intracellular Optical Interrogation with Sub-Cellular Resolution

**DOI:** 10.1101/825489

**Authors:** Menahem Y. Rotenberg, Benayahu Elbaz, Vishnu Nair, Erik Schaumann, Naomi Yamamoto, Laura Matino, Francesca Santoro, Bozhi Tian

**Affiliations:** The James Franck Institute, The University of Chicago, Chicago, Illinois 60637, United States; The Center for Peripheral Neuropathy, The Department of Neurology, The University of Chicago, Chicago, Illinois 60637, United States; Department of Chemistry, The University of Chicago, Chicago, Illinois 60637, United States; The Institute for Biophysical Dynamics, The University of Chicago, Chicago, Illinois 60637, United States; Tissue Electronics, Center for Advanced Biomaterials for Healthcare, Istituto Italiano di Tecnologia, 80125 Naples, Italy; Department of Chemical Materials and Industrial Production Engineering, University of Naples Federico II, 80125 Naples, Italy

## Abstract

Current techniques for intracellular electrical interrogation are substrate bound and are technically demanding, or lack high spatial resolution. In this work, we use silicon nanowires, which are spontaneously internalized by many cell types, to achieve photo-stimulation with sub-cellular resolution. Myofibroblasts loaded with silicon nanowires remain viable and can undergo cell division. Stimulation of silicon nanowires at separate intracellular locations results in local calcium fluxes. We also show that nanowire-containing myofibroblasts can electrically couple to cardiomyocytes in co-culture and that photo-stimulation of the nanowires increases the spontaneous activation rate in neighboring cardiomyocytes. Finally, we demonstrate that this methodology can be extended to the interrogation of signaling in neuron–glia interactions using nanowire-containing oligodendrocytes.

## MAIN TEXT

Intracellular bioelectric interrogation requires the undisruptive introduction of an electrical probe into the cell cytosol. Micropipette electrodes^1, 2^, currently the most widely used tool for intracellular investigations, cannot be used for prolonged times due to mechanical instabilities and the cytosol dilution effect.^3^ Nano-electrode arrays^4, 5^ and field effect transistors^6–8^, powerful new tools for intracellular multiplexed measurements, are substrate-bound and therefore cannot be applied *in vivo* or reconfigured for online recording and stimulation. Optogenetics^9^ presents unique modulation capabilities, allowing optical control at the cellular scale and eliminating the need for direct contact between the light stimulation device and the cell. However, the need for genetic modifications limits the transitional applications of optogenetics, especially to nonhuman primates and other human-relevant models.^10^ Moreover, the basic premise of optogenetics, its elegant utilization of light-activated spatially distributed photosensitive ion channels, limits 3D spatial resolution^11^ and prevents its intracellular application.

Silicon nanowires (SiNWs) offer local optical bioelectric modulation via photo-electrochemical^12^ and photo-thermal^13^ mechanisms. Extracellularly, light-directed stimulation of SiNWs has been used to modulate electrical signals in excitable cells such as neurons^12, 13^ and cardiomyocytes (CMs)^14^. However, SiNWs are also spontaneously internalized by many cell types^15^, leading us to postulate that they may be pre-hybridized with cells to serve as a non-genetic, intracellular, optoelectronic living system. Photo-stimulation using such a system relies on co-localizing high intensity focused light and a SiNW, which also allows for sub-micron spatial resolution in two and three dimensions. In co-culture, the light reflecting properties of SiNWs allow for the identification of the SiNW-containing cells without need for fluorescent genetic labeling.

In this paper, we demonstrate that the localized and cell-specific photo-stimulation produced by this optoelectronic SiNW hybridization cell system enables (i) intracellular interrogation with sub-micron resolution, (ii) cell specific interrogation in heterogenous co-culture, and (iii) hetero-cellular electrical signal transduction investigation (Fig. 1a). We characterize the spontaneous SiNW internalization process in myofibroblasts (MFs), determine that the hybridized cells can undergo cell division, and confirm that the SiNWs remain internalized when the MFs are co-cultured with cardiomyocytes (CMs). We show that photo-stimulation of the internalized SiNWs can be utilized for local intracellular interrogation within the MFs at sub-micron resolution. We demonstrate the ability of the hybridized MFs to electrically couple with co-cultured CMs and report that photo-stimulation of SiNWs within an MF alters the electrical activity of co-cultured CMs. We also establish that this cell-SiNW hybrid system can be extended to the investigation of electrical communication in neuronal systems by demonstrating that photo-stimulation of SiNW-hybridized oligodendrocytes results in calcium flux propagation from the stimulated oligodendrocyte to co-cultured dorsal root ganglion (DRG) neurons.

**Figure 1:**
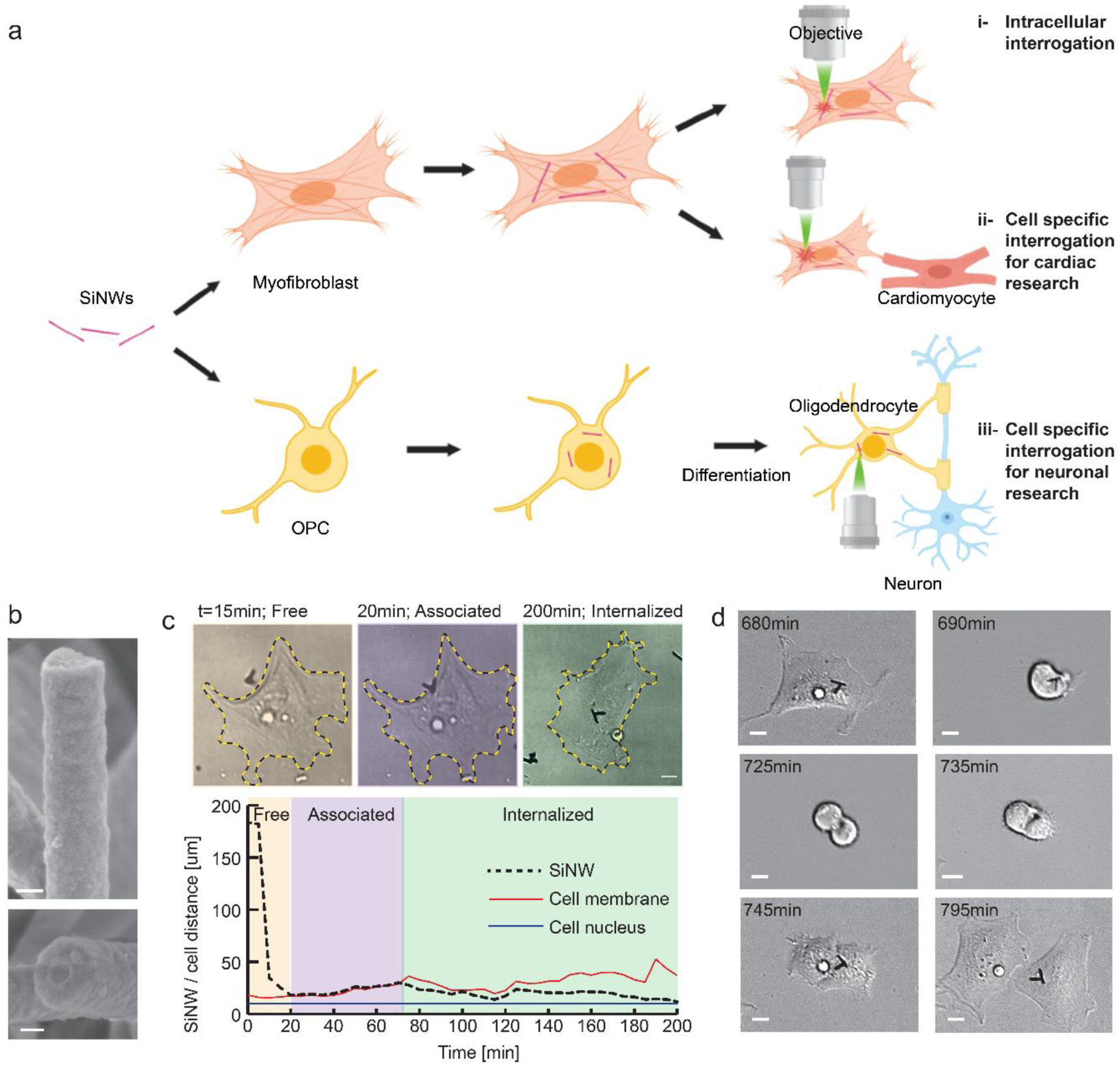
Experimental outline and internalization of the proposed methodology. (a) Silicon nanowires (SiNWs) can be spontaneously internalized by myofibroblasts (MFs) and oligodendrocyte progenitor cells (OPCs). SiNWs can be used for (i) intracellular electrical interrogation with sub-cellular resolution, (ii) cardiac intercellular investigation, or (iii) neuronal intercellular investigation. (b) Scanning electron microscope (SEM) images of the SiNW show one-dimensional morphology and p-i-n core/shell configuration. Scale bars are 100 nm. (c) Top: Phase contrast images of SiNW internalization into MFs, showing free, associated, and internalized SiNWs. Bottom: Time profile of the distance from the center of the nucleus to the SiNW (dashed line), cell membrane (red), and nuclear envelope (blue). (d) Representative phase contrast images of the cell division process. Internalized SiNWs remain within a daughter cell.

Coaxial SiNWs were synthesized as previously reported^12, 16^. Scanning electron microscope (SEM) images show one-dimensional SiNWs with a diameter of ~300 nm and a core-shell semiconducting p-i-n morphology (Fig. 1b). The hybridization process of SiNWs into MFs (Supplementary Video 1) appears to occur in three distinct phases: free SiNWs, associated SiNWs, and internalized SiNWs (Fig. 1c). The time profile of the hybridization process is also shown in Figure 1c. The first section of the plot represents the initial 20 minutes during which the SiNW (dashed line) moves freely in culture media. Upon interaction with the MF cell membrane (red line), free motion stops and the SiNW remains at the border of the cell (the associated phase) for ~50 minutes until the SiNW is internalized. The third section of the plot represents the internalized phase, when the SiNW is in the cytosol between the cell membrane and the nuclear envelope (blue line) (See Supplementary Fig. S1 for more examples). Notably, the SiNW-containing MFs are still capable of undergoing cell division (Fig. 1d and Supplementary Video 1). SiNWs are visible in one of the two daughter cells, suggesting that our hybrid system will be biologically compatible with *in vitro* experimental models.

To confirm that the SiNWs were internalized by the MFs, both confocal and electron microscopy were performed. Following an ultrathin embedding procedure, SEM focused ion beam (SEM-FIB) sectioning of hybridized MF cells was performed.^17^ SEM top view and cross-sections verify that the SiNWs are integrated into the MF cytosol (Fig. 2a). By examining sequential cross-sections, we observe intracellular ultrastructures in tight contact with the surface of the SiNW. These results were corroborated by confocal imaging using MF-SiNW hybrids labeled with cytosolic and membranal markers (Fig. 2b, and Supplementary Fig. S2). Cross-sectional z-stacks of the confocal images show that the SiNWs are contained within the cytosol of MFs.

**Figure 2:**
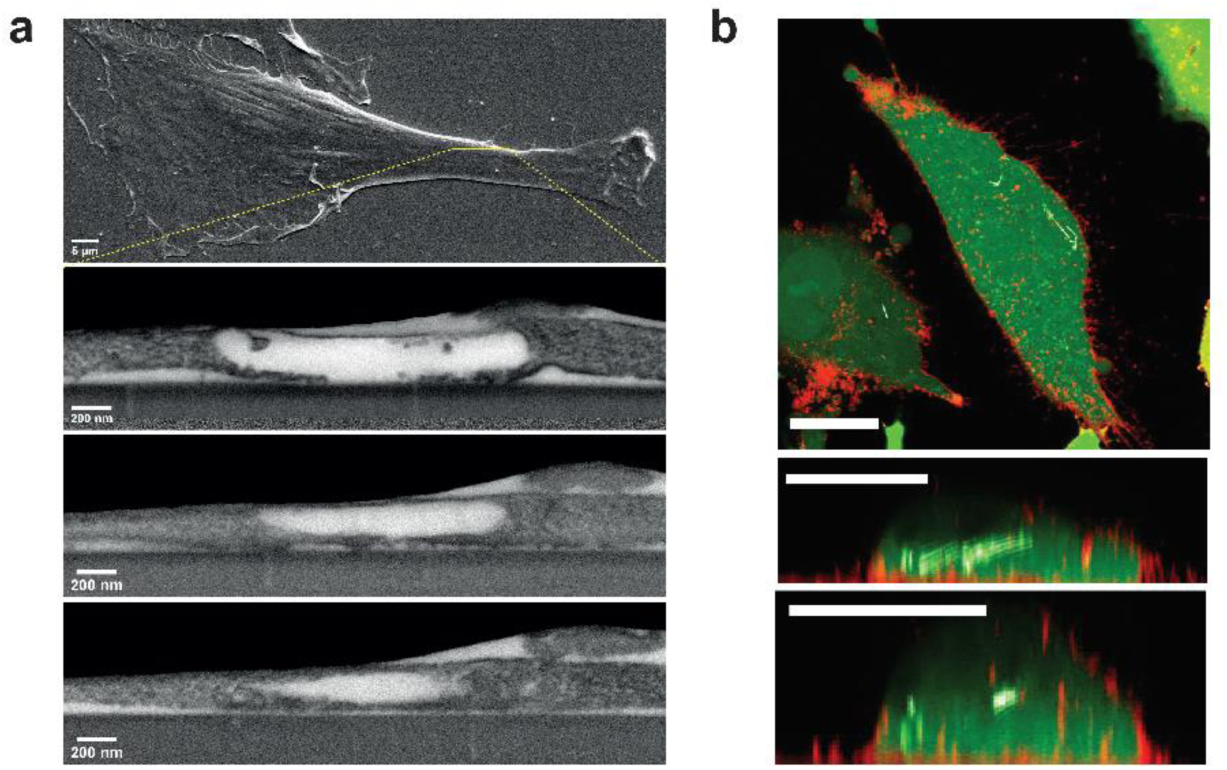
The intracellular SiNW-MF interface. (a) Scanning electron and focused ion beam microscopy (SEM-FIB) images of three cross-sections showing fully internalized SiNWs. (b) Confocal microscopy images of cytosolic (green) and membranal (orange) markers show fully internalized SiNWs (white) in the cytosol of the MF. Scale bars are 20 µm.

Our SiNW-MF hybrids are unique tools for interrogating bioelectric dynamics from different sub-cellular regions within a single cell. To demonstrate this capability, we sequentially photo-stimulated different SiNWs in a single MF-SiNW hybrid cell with a single 4 mW, 1 ms laser pulse (Fig. 3a-b and Supplementary Video 2). Optical mapping reveals a brief and reversible calcium flux originating from the stimulation target as a result of the photo-stimulation (Fig. 3b, panels 1-3), demonstrating that the SiNW offers spatial control over calcium dynamics. This result is also observed after multiple stimulations of the same cell, indicating that the cell remains intact after the 4 mW laser pulses. Increased laser power of 7 mW (Fig. 3b, panel 4) induces a larger calcium flux over the same time scale. These intensity-dependent results suggest that the mechanism of MF electrical activity differs from that of CMs and other excitable cells. Whereas excitable cells operate on a binary fire/no fire action potential (AP), MFs appear to display stimulation intensity-dependent calcium flux and propagation. To better demonstrate this intensity dependence, Figure 3c shows kymographs corresponding to the stimulation in Figure 3b. Each kymograph demonstrates the intensity dependent response to a particular response in terms of velocity curve (v) and distance of propagation (d).

**Figure 3:**
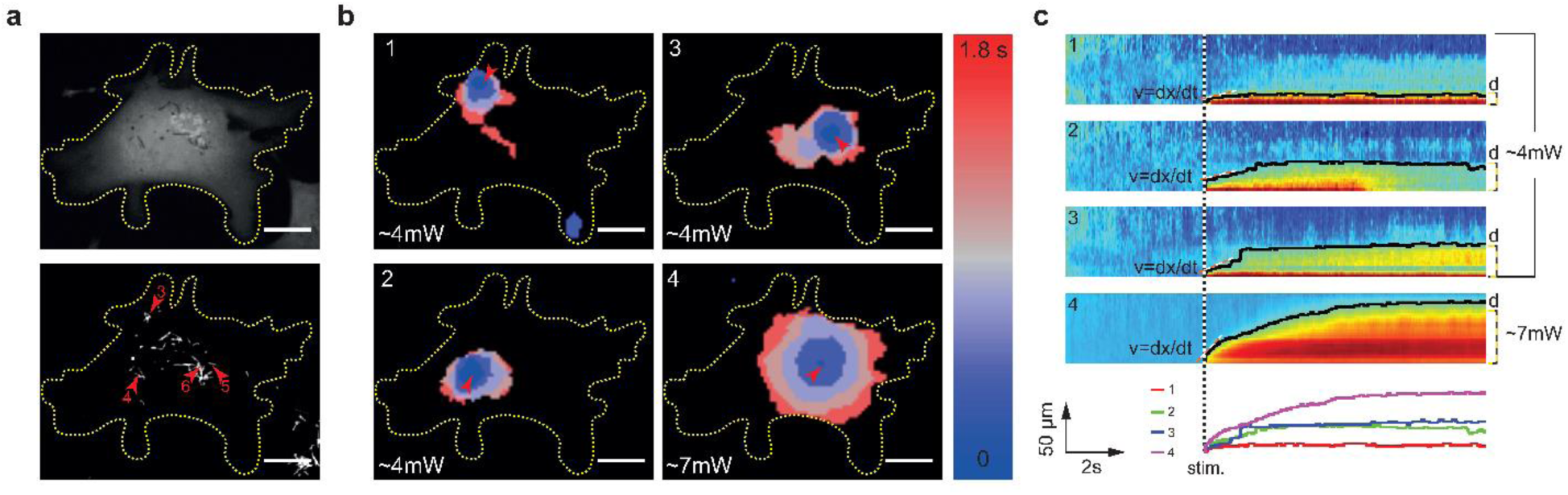
Intracellular electrical interrogation with sub-cellular resolution. (a) Fluorescent (fluo-4; top) and dark field (bottom) images of SiNW-containing MF. Red arrowheads indicate stimulated SiNWs. (b) Heatmaps show calcium propagation within the MF-SiNW hybrid at different stimulation locations at laser power of 4 mW (panels 1-3) and 7 mW (panel 4). (c) Kymograph of dF/F represents the calcium flux from the stimulated SiNW. Panels 1-4 correspond to stimulation 1-4 from Figure 3(b). More rapid calcium propagation occurs at higher laser power, as shown by the dF/F profile over time (bottom).

To utilize the SiNW-hybrid system for interrogation of intercellular electrical signal communication between MFs and CMs, the SiNWs must remain within the MFs during co-culture with CMs. To confirm this retention, we performed immunocytochemical imaging of CMs co-cultured with SiNW-containing MFs. As seen in Figure 4a, SiNWs are located almost exclusively in MFs (red), not in CMs (green) (see also Supplementary Fig. S3). In general, we found only two isolated cases in which a SiNW associated with a CM. In both cases, only a single SiNW was involved and its location (intra- or extra-cellular) was inconclusive. Statistical analysis of the number of SiNWs associated with MFs and CMs (Fig. 4b, top) and the fraction of hybridized cells (those with at least one internalized SiNW; Fig 4b, bottom) further demonstrate that SiNWs remained inside the MFs.

**Figure 4:**
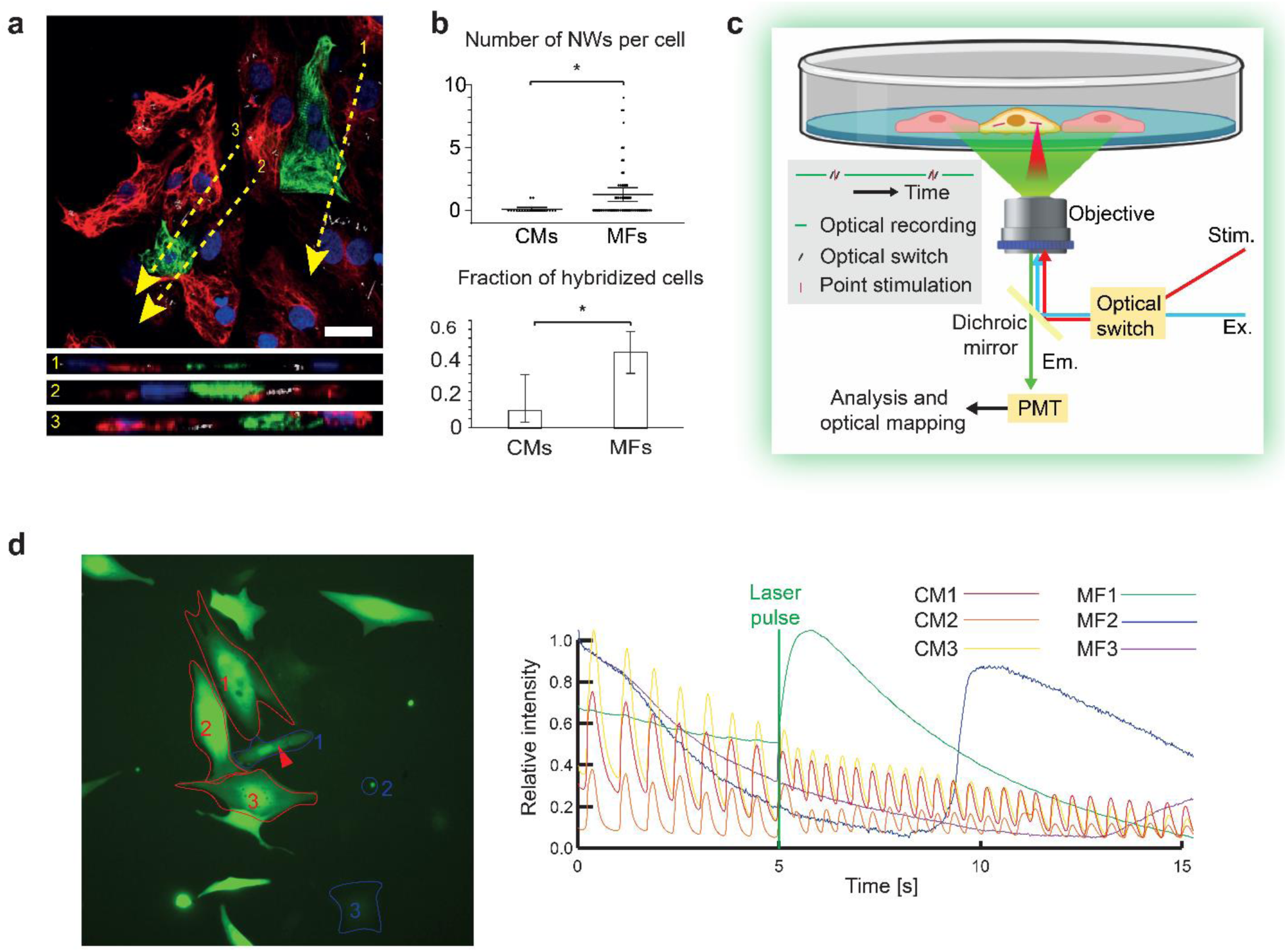
Cell-specific stimulation in cardiac cells co-culture. (a) Immunofluorescence confocal images show that SiNWs (white) are internalized exclusively by MFs (red) when co-cultured with CMs (green). Yellow arrows represent cross-sectioned z-stacks (bottom). Scale bar is 20 µm. (b) Statistical analysis of the number of SiNWs within each MF and CM (top, p < 0.0001) and the fraction of cells hybridized with at least one SiNW out of the general cell population (bottom, p < 0.05). Error bars represent 95% confidence interval of the data from 20 CMs and 62 MFs. (c) SiNWs are stimulated by a laser. Changes in calcium levels are imaged using calcium sensitive dye (fluo-4) and fluorescent microscopy. (d) Left: Fluorescent image of MF-SiNW/CM co-culture treated with Fluo-4. Regions of interest are indicated in red (CMs) and blue (MFs) Right: dF/F intensity versus time profile for regions of interest before and after stimulation of MF 1 (1ms, 7 mW). The immediate increase of CM activation rate following stimulation is evident.

To verify that the hybrids are able to electrically couple to other cells, we optically stimulated MF-SiNW hybrids in co-culture with CMs and visualized the response in the CMs (Fig. 4c). Plotting the relative intensity profile versus time of the different cells prior to the optical stimulation, we identify the CMs in the co-culture as they were spontaneously beating and fully synchronized. The MFs in the co-culture are static with no apparent change in calcium levels (Fig. 4d). Optical stimulation of an MF-SiNW hybrid (red arrow) results in an immediate calcium flux within the hybrid cell (Fig. 4d; green line in plot). We attribute this increase in calcium level to a combination of photo-thermal (*i.e.*, producing transient membrane poration) and photo-electrochemical (*i.e.*, producing ROS) processes. As a result of stimulation, the contraction rate of the associated CMs immediately increases from ~1.4 Hz to ~2.6 Hz (Fig. 4d), indicating that stimulation of the MF-SiNW hybrid modifies the electrical activity of neighboring CMs. This frequency increase may be attributed to an increase in the CM resting membrane potential resulting from the intracellular calcium increase within the coupled MFs. Interestingly, the calcium flux in the stimulated MF (MF1 in Fig. 4d) slowly propagated to the neighboring MFs (MF2 and MF3), as illustrated by their sequential florescence increase. These findings are supported by previous work which showed that MF propagation of electrical activity *in vitro* is characterized by local conduction delays.^18^

To demonstrate the flexibility of our cell-SiNW system and its application to other cell types, we expanded our methodology to neuron-glia interactions. Calcium transients influence the myelination of neuronal axons by oligodendrocyte lineage cells,^19–24^ but the degree to which these transients are induced by neuronal activity^19–22^ or by oligodendrocytes themselves^23, 24^ is unclear^25^. As a proof-of-concept that our cell-SiNW hybrids may be used as a new tool in the investigation of calcium transients in neuron-glia interactions, we hybridized SiNWs with oligodendrocyte progenitor cells (OPCs). We co-cultured the SiNW-OPC hybrids with DRG neurons in differentiation medium, which induced the OPC cells to differentiate into oligodendrocytes. Interactions between oligodendrocytes and DRG neurons are visible in Figure 5a. Co-localization of myelin basic protein (red) and neuronal marker NueN (green) represents sections of neuronal axons that have been myelinated by an associated oligodendrocyte. Upon optical stimulation of the SiNW within the oligodendrocyte (red arrowhead, Fig. 5b), a calcium flux propagated from the SiNW in the cell soma to the associated neuron (Fig. 5b, [bottom] and Supplementary Video 3). Although optical mapping appears to show a gap in the propagation through the oligodendrocyte soma, closer examination of that region by intensity profiles reveals that the calcium indeed propagates through the cell process. However, given that the size of the cell process is decreasing away from the soma, the fluorescent intensity in these regions is under the threshold of the optical mapping algorithm (Supplementary Fig. S4).

**Figure 5:**
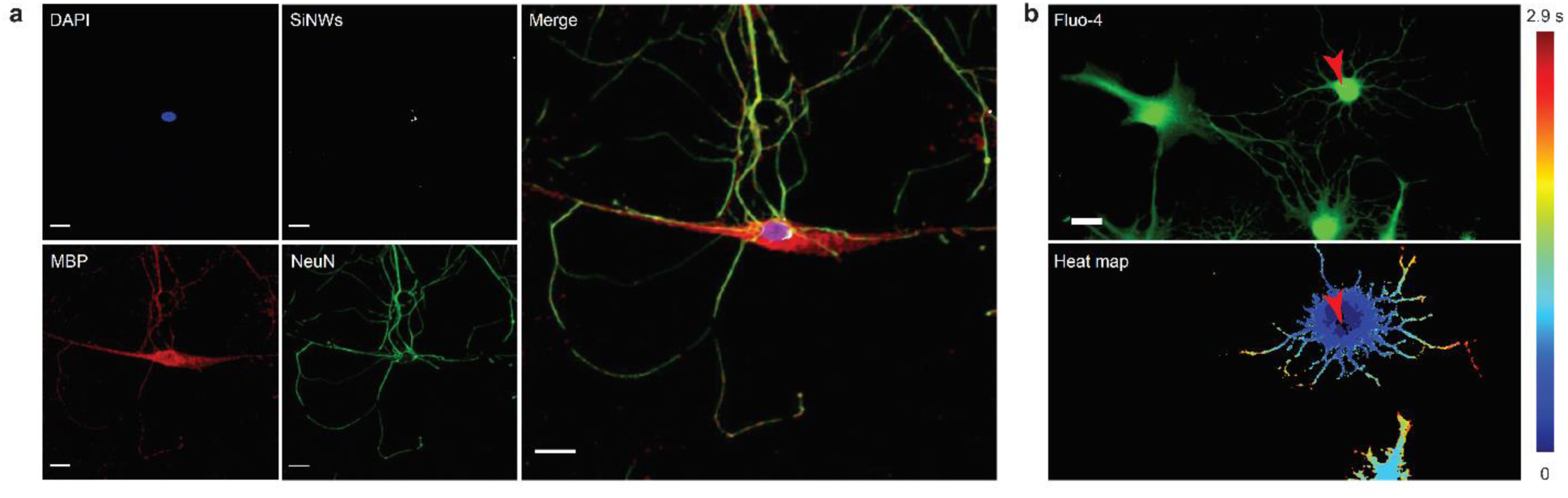
Cell-specific stimulation for investigating neuron-glia interactions. (a) Representative immunofluorescence confocal images of an oligodendrocyte (MBP, red) myelinating a DRG neuron (NeuN, green). Internalized SiNWs (white) can be seen near the MF nucleus (DAPI, blue). (b) Top: Fluorescent image of oligodendrocyte/DRG neuron co-culture treated with Fluo-4. Bottom: Heatmap of the calcium flux propagating from the oligodendrocyte to the DRG neuron upon SiNW photo-stimulation at the red arrowhead.

In summary, we have characterized and demonstrated the utility of a new SiNW-based tool for intracellular electrical interrogation. The process of SiNW hybridization with different cells is elegant in its simplicity, relying on spontaneous hybridization rather than abrasive electroporation or sonication. Initiation of intracellular stimulation using the SiNW hybrids does not require physical access to the cytosol or genetic modifications. The SiNW hybrids also provide sub-micron resolution with standard microscopy techniques. Internalized SiNWs clearly distinguish the hybridized cells from other cells in co-culture, eliminating the need for genetic or fluorescent labeling to identify cell types. Because the hybridized cells retain their ability to proliferate and electrically couple with other cells, we anticipate that this approach will expand the available tool kit for bioelectronic studies. Our work has demonstrated the versatility of these hybrids in *in vitro* cardiac and neural settings, but later work may expand our understanding of complex, multi-cellular electrical cascades in *in vivo* studies.

## ASSOCIATED CONTENT

The Supporting Information is available free of charge on the ACS Publications website at DOI -----:

Additional experimental details and SI figures S1-S4 with examples of the internalization process and confocal images (PDF)

SiNW internalization and cell division (MOV)

Intracellular calcium wave propagation (dF/F) within a MF at subcellular resolution (MOV)

Intercellular calcium wave propagation (dF/F) from an oligodendrocyte to a DRG neuron (MOV)

## AUTHOR INFORMATION

### Author Contributions

The manuscript was written through contributions of all authors. All authors have given approval to the final version of the manuscript.

### Funding Sources

This work is supported by the Air Force Office of Scientific Research (AFOSR FA9550-18-1-0503), US Army Research Office (W911NF-18-1-0042), US Office of Naval Research (N000141612530, N000141612958), and the National Institutes of Health (NIH NS101488). This work was partially supported by the NIH (R01NS067550) to B.E.

## ACKNOWLEDGMENTS

We thank Dr. Karen Watters for scientific editing of the manuscript. We would like to thank the Popko Lab for their helpful comments. We thank Professor Margaret Gardel from the University of Chicago for her technical assistance. We thank the Integrated Light microscopy Core Facility at the University of Chicago for their technical assistance.

## REFERENCES

1. Sakmann, B.; Neher, E. Annu Rev Physiol 1984, 46, (1), 455–72.

2. Jayant, K.; Hirtz, J. J.; Plante, I. J.; Tsai, D. M.; De Boer, W. D.; Semonche, A.; Peterka, D. S.; Owen, J. S.; Sahin, O.; Shepard, K. L.; Yuste, R. Nat Nanotechnol 2017, 12, (4), 335–342.

3. Molleman, A., Patch clamping: an introductory guide to patch clamp electrophysiology. John Wiley & Sons: 2003.

4. Xie, C.; Lin, Z.; Hanson, L.; Cui, Y.; Cui, B. Nat Nanotechnol 2012, 7, (3), 185–90.

5. Robinson, J. T.; Jorgolli, M.; Shalek, A. K.; Yoon, M. H.; Gertner, R. S.; Park, H. Nat Nanotechnol 2012, 7, (3), 180–4.

6. Duan, X.; Gao, R.; Xie, P.; Cohen-Karni, T.; Qing, Q.; Choe, H. S.; Tian, B.; Jiang, X.; Lieber, C. M. Nat Nanotechnol 2011, 7, (3), 174–9.

7. Tian, B.; Cohen-Karni, T.; Qing, Q.; Duan, X.; Xie, P.; Lieber, C. M. Science 2010, 329, (5993), 830–4.

8. Qing, Q.; Jiang, Z.; Xu, L.; Gao, R.; Mai, L.; Lieber, C. M. Nat Nanotechnol 2014, 9, (2), 142–7.

9. Fenno, L.; Yizhar, O.; Deisseroth, K. Annual review of neuroscience 2011, 34.

10. Galvan, A.; Stauffer, W. R.; Acker, L.; El-Shamayleh, Y.; Inoue, K.-i.; Ohayon, S.; Schmid, M. C. Journal of Neuroscience 2017, 37, (45), 10894–10903.

11. Packer, A. M.; Russell, L. E.; Dalgleish, H. W.; Hausser, M. Nat Methods 2015, 12, (2), 140–6.

12. Parameswaran, R.; Carvalho-de-Souza, J. L.; Jiang, Y.; Burke, M. J.; Zimmerman, J. F.; Koehler, K.; Phillips, A. W.; Yi, J.; Adams, E. J.; Bezanilla, F.; Tian, B. Nat Nanotechnol 2018, 13, (3), 260–266.

13. Fang, Y.; Jiang, Y.; Acaron Ledesma, H.; Yi, J.; Gao, X.; Weiss, D. E.; Shi, F.; Tian, B. Nano letters 2018, 18, (7), 4487–4492.

14. Parameswaran, R.; Koehler, K.; Rotenberg, M. Y.; Burke, M. J.; Kim, J.; Jeong, K.-Y.; Hissa, B.; Paul, M. D.; Moreno, K.; Sarma, N. Proceedings of the National Academy of Sciences 2019, 116, (2), 413–421.

15. Zimmerman, J. F.; Parameswaran, R.; Murray, G.; Wang, Y.; Burke, M.; Tian, B. Sci Adv 2016, 2, (12), e1601039.

16. Tian, B.; Zheng, X.; Kempa, T. J.; Fang, Y.; Yu, N.; Yu, G.; Huang, J.; Lieber, C. M. nature 2007, 449, (7164), 885.

17. Li, X.; Matino, L.; Zhang, W.; Klausen, L.; McGuire, A. F.; Lubrano, C.; Zhao, W.; Santoro, F.; Cui, B. Nat Protoc 2019, 14, (6), 1772–1802.

18. Gaudesius, G.; Miragoli, M.; Thomas, S. P.; Rohr, S. Circ Res 2003, 93, (5), 421–8.

19. Krasnow, A. M.; Ford, M. C.; Valdivia, L. E.; Wilson, S. W.; Attwell, D. Nat Neurosci 2018, 21, (1), 24–28.

20. Wake, H.; Lee, P. R.; Fields, R. D. Science 2011, 333, (6049), 1647–51.

21. Friess, M.; Hammann, J.; Unichenko, P.; Luhmann, H. J.; White, R.; Kirischuk, S. Cell Calcium 2016, 60, (5), 322–330.

22. Hines, J. H.; Ravanelli, A. M.; Schwindt, R.; Scott, E. K.; Appel, B. Nat Neurosci 2015, 18, (5), 683–9.

23. Baraban, M.; Koudelka, S.; Lyons, D. A. Nat Neurosci 2018, 21, (1), 19–23.

24. Battefeld, A.; Popovic, M. A.; de Vries, S. I.; Kole, M. H. P. Cell Rep 2019, 26, (1), 182–191 e5.

25. Elbaz, B.; Popko, B. Trends Neurosci 2019.

