## Supplementary information for "Silicon Nanowires for Intracellular Optical Interrogation with Sub-Cellular Resolution"

### Methods:

**Nanowire synthesis.** SiNWs, with p-i-n core-shell configuration, were synthesized using an Au nanocluster-catalyzed chemical vapor deposition (CVD) process. Au colloidal nanoparticles (Ted Pella, 100 nm diameter) were suspended in 5% hydrofluoric acid and deposited onto Si substrates (Nova Electronic Materials). SiNWs were grown using silane ($SiH_4$) as the Si reactant and diboron ($B_2H_6$, 100 ppm in $H_2$) as the p-type core dopant. Phosphine ($PH_3$, 1000 ppm in $H_2$) was used as the n-type shell dopant, and hydrogen ($H_2$) as the carrier gas. For p-type core growth, $SiH_4$, $B_2H_6$, and $H_2$ were delivered at flow rates of 2, 10, and 60 standard cubic centimeters per min (sccm), respectively. Growth was carried out for 30 min at 470°C and 40 torr. Then, intrinsic Si shell (i-shell) deposition was performed. The temperature was ramped up to 650°C with no gas flow under vacuum. Then, $SiH_4$ and $H_2$ were delivered at 0.3 and 60 sccm, respectively, for 20 min at 15 torr. For the n-type outer shell, $PH_3$ gas was added at a flow rate of 1.5 sccm, under the same conditions, for 20 min.

**Cardiac cell culture**. All animal procedures were approved by the University of Chicago Institutional Animal Care and Use Committee (IACUC) and conducted in complete compliance with the IACUC Animal Care and Use Protocol. Hearts were excised from P0-5 neonatal rats into ice cold Hanks' Balanced Salt solution (HBSS) without $Ca^{2+}$ or $Mg^{2+}$. Primary cardiomyocytes (CMs) and cardiac fibroblasts were isolated using the Pierce™ Primary Cardiomyocyte Isolation Kit (Thermo Fisher Scientific). Procedure followed manufacturer's protocol. To separate fibroblasts from CMs, the suspended cells were pre-plated for 1-2 hr, allowing the fibroblasts to adhere to the tissue culture plate. The enriched CM floating suspension was seeded on fibronectin (Sigma) treated glass bottom dishes. Fibroblasts were cultured (DMEM high glucose + 10% FBS, 1% Glutamax and 1% penicillin–streptomycin) and proliferated until cells reached ~80% confluency. Proliferating fibroblasts were considered to be myofibroblasts (MFs) as they spontaneously differentiate in standard culture[1]. MFs were used for hybridization with SiNWs and co-culture with CMs. For seeding SiNWs, 3 × 3 $mm^2$ of substrate with CVD-grown SiNWs was sonicated for 10 min in culture media and then seeded on a 100 mm tissue culture dish (~55 $cm^2$, Falcon). After 12 hr, cells were vigorously rinsed five times until free-floating SiNWs were eliminated. MF-SiNW hybrids were harvested using trypsin for 2 min at 37°C and then rinsed and centrifuged for 5 min at 200g. The harvested MF-SiNW hybrid cells were reseeded alone or co-cultured with CMs.

**Live imaging of the SiNW internalization process.** Collagen-coated glass bottom dishes were pre-plated with MFs for 10 min to avoid CM contamination. The cells were allowed to proliferate for 24 hr. SiNWs were seeded onto MFs (keeping the same ratio of 9 $mm^2$ of Si substrate with CVD-grown SiNWs to ~55 $cm^2$ culture dish) immediately before live imaging began. Imaging was performed with a multimode high-resolution epi-fluorescence microscope (Nikon) using phase contrast. Images were taken every 5 min.

**Scanning Electron and Focused Ion Beam Microscopy (SEM-FIB).** MF-SiNW hybrid cells were seeded on glass cover slips and then processed following the ultra-thin plasticization (UTP) protocol[2]. Cells were fixed with 2.5% glutaraldehyde (Sigma-Aldrich) diluted in 0.1 M sodium cacodylate (Electron Microscopy Sciences) for 1 hour at room temperature and then rinsed with 0.1 M sodium cacodylate buffer. Afterwards, cells were rinsed in 20 mM glycine (Sigma-Aldrich) to quench unreacted aldehydes. Cells were post-fixed in 2% osmium tetroxide and 2% potassium ferricyanide (both Electron Microscopy Sciences) for 1 hr at 4°C in the dark. After washing, cells were incubated with 1% thiocarbohydrazide (Sigma-Aldrich), and immersed in 2% osmium tetroxide. Afterwards, cells were left overnight in 4% uranyl acetate (Sigma-Aldrich) at 4°C and rinsed with deionized water at 4°C. Cells were treated for 3 min with 0.15% tannic acid

(Sigma-Aldrich) and dehydrated with ethanol (30%, 50%, 70%, 90%, 100%, Sigma-Aldrich). Absolute ethanol was gradually replaced with Spurr's low viscosity embedding resin mixture (Electron Microscopy Sciences) to preserve cell structures. Embedding was performed by gradually increasing the resin-ethanol ratio. Minimal resin covering of cells was achieved by upright positioning of the cell substrate for 2 hr and quickly rinsing with absolute ethanol. Cells were polymerized in an oven overnight and then mounted with conductive silver paste (RS PRO) onto a pin stub.

**Confocal images of internalized SiNWs**. MF-SiNW hybrids were harvested, seeded on glass bottom dishes and allowed to sit for 24 hr. Cells were loaded with live dye (Calcein, AM, cell-permeant dye - Thermo Fisher Scientific) and membrane marker (CellMask Orange Plasma membrane Stain, Thermo Fisher Scientific). The cells were then immediately imaged using a Leica SP5 Tandem Scanner Spectral 2-Photon Confocal. Z-stacks were cross-sectioned and processed using Fiji software[3].

**Dorsal root ganglion (DRG) neurons and oligodendrocytes isolation.** All animal procedures were approved by the University of Chicago Institutional Animal Care and Use Committee (IACUC) and conducted in complete compliance with the IACUC Animal Care and Use Protocol. For DRG neurons, p0-3 neonatal rats were decapitated, and their spinal column cut open along the middle. The ganglions were isolated from the spinal column using a fine tip forceps and placed in ice-cold DMEM/F12 (Invitrogen). Ganglia were digested in 2.5 mg/mL of of Trypsin (Worthington) in EBSS without $Ca^{2+}$ or $Mg^{2+}$ for 20 min. They were transferred into EBSS with 10% FBS and triturated with fire polished glass pipette in decreasing order or diameter to ensure homogeneous separation. Cells were resuspended in complete media (DMEM/F12 with 1% pen-strep, 5 % FBS along with 20 µM 5-fluoro uracil, Sigma) and 50 ng/mL of NGF 2.5S (Thermofisher invitrogen)). After 24 hours, the complete media was supplemented with 40 µM uridine (Sigma Aldrich) and 1 µM Arabinofuranosyl Cytidine (Sigma). DRG neurons were seeded on glass bottom dishes treated with poly-D lysine.

Mouse oligodendrocyte progenitor cells were isolated from 6 day-old pups by immunopanning as we previously described[4]. Mouse oligodendrocyte progenitor cells were maintained in proliferation media containing PDGFRa. One day before seeding the cells on the DRGs, the oligodendrocyte progenitor cells were incubated for 4 hours with SiNWs, and then washed with the same medium. The SiNW-loaded oligodendrocyte progenitor cells were seeded on DRG neurons and allowed to differentiate for 4 days in differentiation medium containing T3.

**Calcium sensitive dye**. Cells were treated with calcium sensitive dye (2 µM Fluo-4, AM, cell permeant, Thermo Fisher Scientific) for 30 min at 37°C. Then, cells were rinsed and internalized fluo-4 was allowed to undergo de-esterification for 30 more min at 37C. The treated cells were visualized with different microscopes: (i) A Marianas Yokogawa type spinning disk confocal was used for visualizing and stimulating the cells in Fig. 3, (ii) A Nikon TI2-E inverted microscope with a Photosim Scanner for Fig. 4d. and (iii) A Leica SP5, STED-CW Superresolution Laser Scanning Confocal for Fig. 5b.

**Immunocytochemistry**. Cells were fixed with 4% paraformaldehyde for 10 min (Sigma) and permeabilized (0.2% Triton X-100, Sigma). The cells were blocked (2% BSA) to prevent non-specific binding. Then, cardiac cells were incubated with anti-rabbit cardiac troponin I polyclonal antibody (Abcam; for CMs) and anti-chicken vimentin polyclonal antibody (Abcam; for MFs) and oligodendrocytes/DRG co-culture were incubated with anti-rabbit NeuN polyclonal antibody (Abcam; for DRG neurons) and anti-mouse myelin basic protein (MBP) polyclonal antibody (Abcam; for oligodendrocytes). Cells were then incubated with Alexa Fluor 488 and Alexa Fluor 647 secondary antibodies (Abcam). ProLong Gold Antifade Mountant with

DAPI (Themo Fisher) was used to label the nuclei. Cells were imaged using the Leica SP5 Tandem Scanner Spectral 2-Photon Confocal.

**Optical mapping**. Calcium sensitive dye videos were analyzed using ImageJ[5] and an online available macro[6] for ΔF/F movies. A threshold was manually selected to convert the videos to a binary activated/not activated video, and the resulting stack was used to generate a time color code for the activation propagation. Kymographs were made by cross sectioning a fixed length of the ΔF/F movies, originating at the stimulation site. The x-axis is time and the y-axis is the distance along the cross-section. The boundary lines were created by thresholding the kymograph, while the same thresholding values were used for all videos.

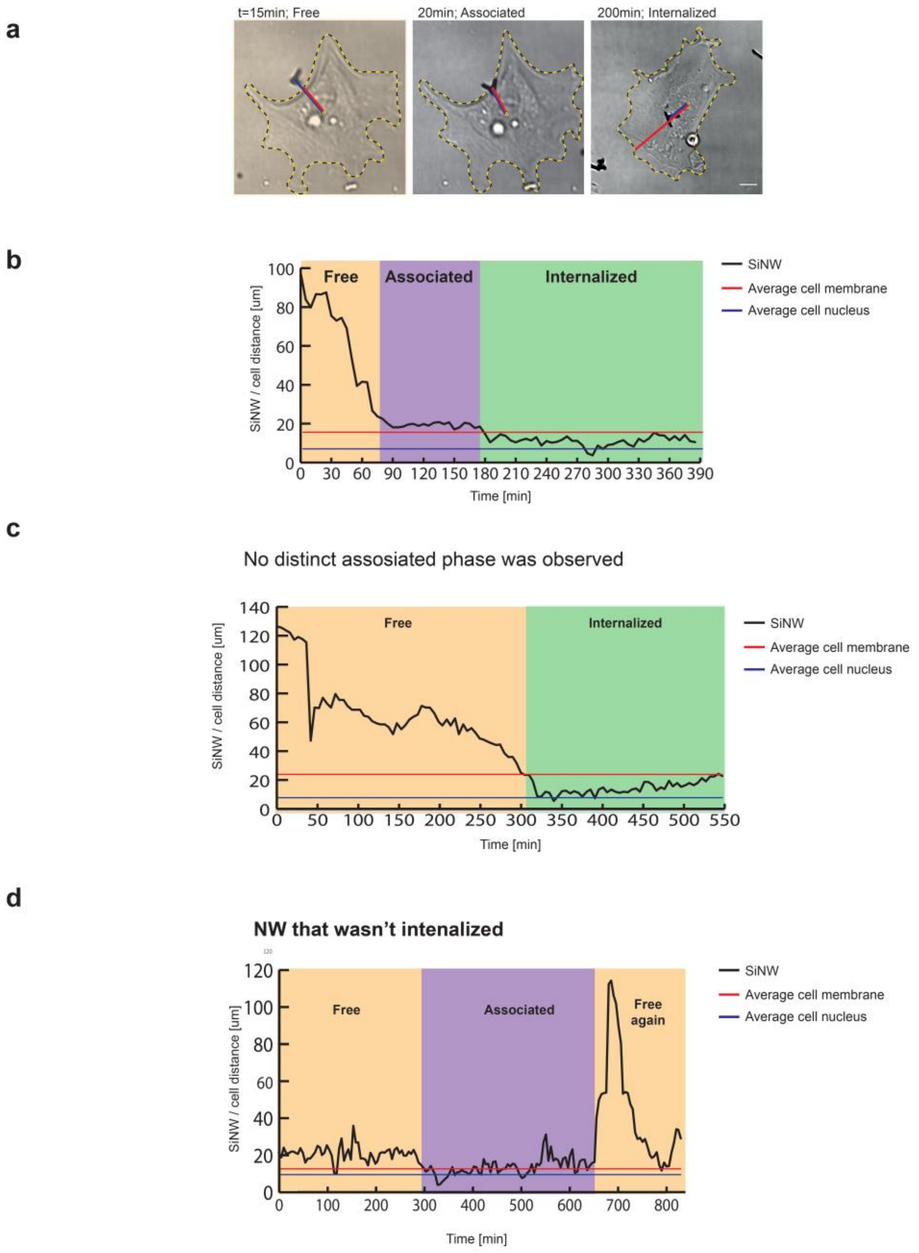


Supplemental Figure 1: Further detail and examples of the SiNW internalization process and analysis. (a) Representative images from Fig. 1C showing the measurement of distance from the nucleus to the SiNW (blue) and the plasma membrane (red). See methods for further details. (b) Additional time profile example of the three distinct phases of SiNW internalization. Average cell radius (red line) and average nucleus radius (blue line) are used for clarity. (c) An example of the hybridization process in which an immediate internalization of the SiNW had occurred without an associated phase. Average cell radius (red line) and average nucleus radius (blue line) are used for clarity. (d). An example of a SiNW that was not internalized by the cell. Such SiNWs are removed by the wash steps prior to co-culture (see methods). Average cell radius (red line) and average nucleus radius (blue line) are used for clarity.

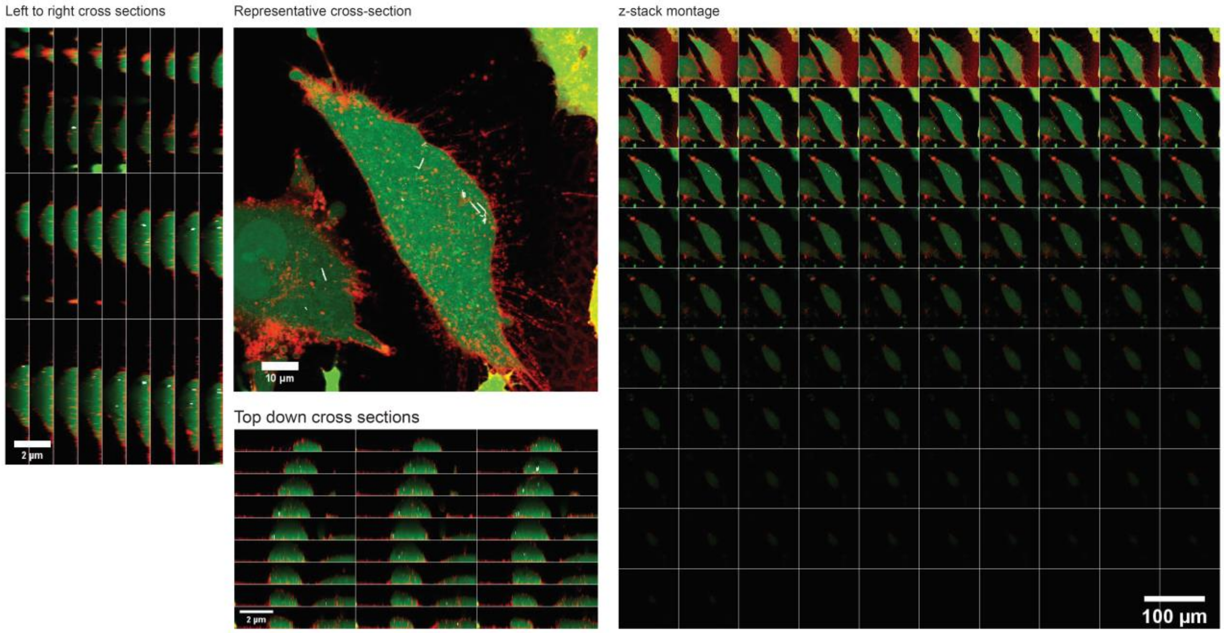


Supplemental Figure 2: Z-stack and cross-sectional view of confocal imaging of the SiNW-loaded MF. Cytoplasm (Cacein AM, green), plasma membrane (CellMask, orange), and SiNWs (reflected light, white) are all visible. A representative z cross-section shows cell volume (top center). Individual z-stack images (right), y-z cross sections (left), and x-z cross sections (bottom center) are also shown.

Example 1:

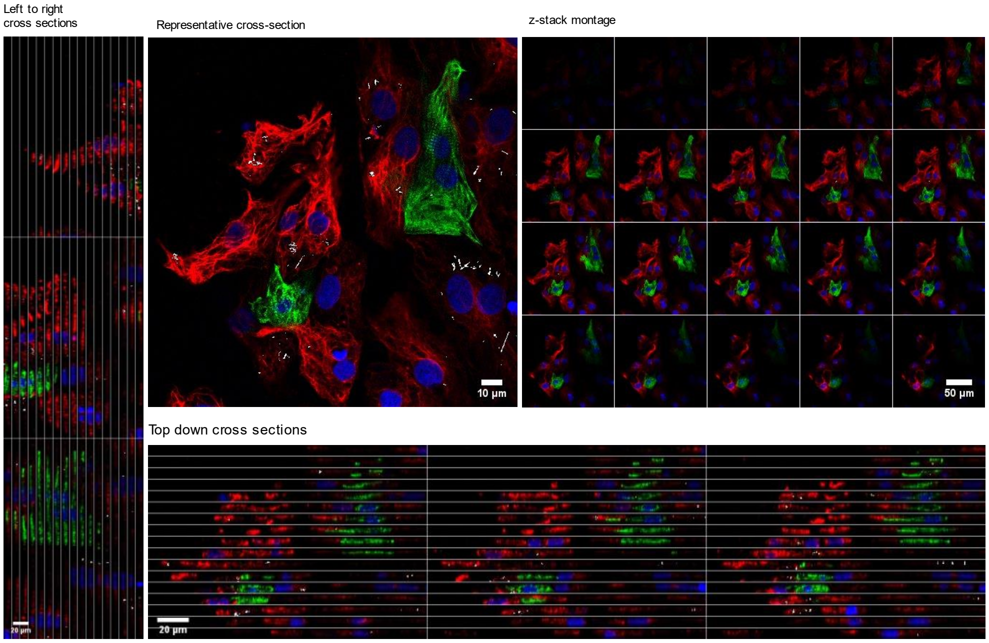


Example 2:

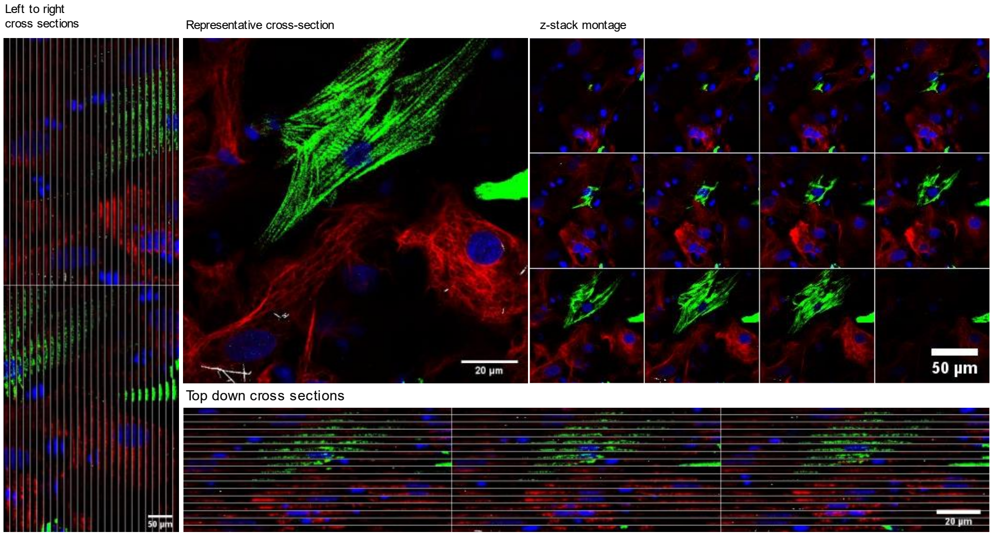


Supplemental Figure 3: Confocal imaging of MF-CM co-cultures. Two examples of the co-culture show the distribution of SiNW within the cells. MFs are identified by vimentin staining (red), CMs are identified by cardiac troponin staining (green), and SiNWs are visible by reflected light (white). Nuclei are also stained with DAPI (blue). A representative z cross section, y-z cross section, x-z cross section, and mosaics of whole z-stacks are also presented. The distribution illustrates that the SiNWs are almost entirely associated with MFs.

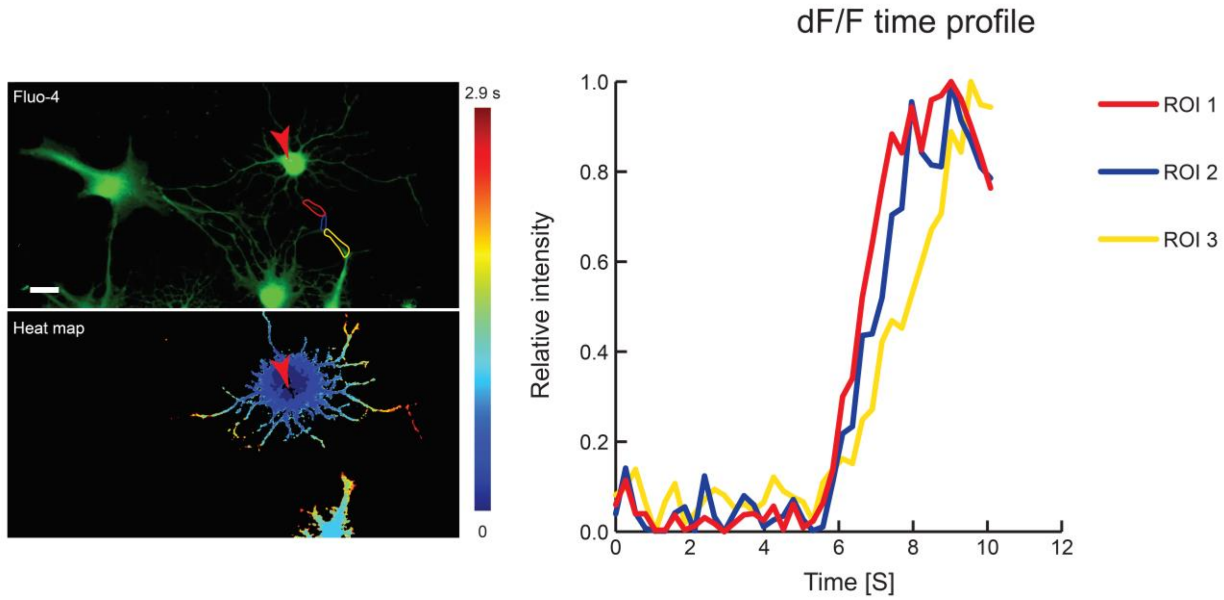


Supplemental Figure 4: Calcium propagation between oligodendrocytes and DRG neurons. Three regions of interest (top left, shown in red, blue, and yellow) are selected to demonstrate the propagation through the oligodendrocyte. Time profile of dF/F (right) displays activation timing indicating sequential flux towards the neuron, despite flux being undetectable by heat map (bottom left).

Supplemental Video 1:

Phase contrast imaging of the internalization process. Images were acquired every 5 minutes. Different phases of the process are highlighted in the video. After internalization is complete, the MF-SiNW hybrid undergoes cell division.

Supplemental Video 2:

Intracellular interrogation with sub-cellular resolution is presented by the dF/F videos of four consecutive stimulations of the same MF-SiNW hybrid. Location of stimulation is marked by red arrowheads and laser power is denoted. Numbers in the video correspond to the numbering of the heatmap in Fig. 3a.

Supplemental Video 3:

Intercellular calcium propagation from the stimulated oligodendrocyte to the DRG neuron is shown via dF/F videos following optical stimulation.


1 Rohr, S. Cardiac fibroblasts in cell culture systems: myofibroblasts all along? *Journal of cardiovascular pharmacology* **57**, 389-399 (2011).

2 Santoro, F. *et al.* Revealing the cell–material interface with nanometer resolution by focused ion beam/scanning electron microscopy. *ACS nano* **11**, 8320-8328 (2017).

3 Schindelin, J. *et al.* Fiji: an open-source platform for biological-image analysis. *Nat Methods* **9**, 676-682, doi:10.1038/nmeth.2019 (2012).

4 Elbaz, B. *et al.* Phosphorylation state of ZFP24 controls oligodendrocyte differentiation. *Cell reports* **23**, 2254-2263 (2018).

5 Rueden, C. T. *et al.* ImageJ2: ImageJ for the next generation of scientific image data. *BMC Bioinformatics* **18**, 529, doi:10.1186/s12859-017-1934-z (2017).

6 Ackman, J. *dFoFmovie-CatFullAutoSave.java*, <https://gist.github.com/ackman678/11155761> (no date).
